# A standard protocol to report discrete stage-structured demographic information

**DOI:** 10.1101/2023.01.13.523871

**Authors:** Samuel J. L. Gascoigne, Simon Rolph, Daisy Sankey, Nagalakshmi Nidadavolu, Adrian S. Stell Pičman, Christina M. Hernández, Matthew E. R. Philpott, Aiyla Salam, Connor Bernard, Erola Fenollosa, Young Jun Lee, Jessie McLean, Shathuki Hetti Achchige Perera, Oliver G. Spacey, Maja Kajin, Anna C. Vinton, C. Ruth Archer, Jean H. Burns, Danielle L. Buss, Hal Caswell, Judy P. Che-Castaldo, Dylan Z. Childs, Pol Capdevila, Aldo Compagnoni, Elizabeth Crone, Thomas H. G. Ezard, Dave Hodgson, Tiffany M. Knight, Owen R. Jones, Eelke Jongejans, Jenni McDonald, Brigitte Tenhumberg, Chelsea C. Thomas, Andrew J. Tyre, Satu Ramula, Iain Stott, Raymond L. Tremblay, Phil Wilson, James W. Vaupel, Roberto Salguero-Gómez

**Affiliations:** Department of Biology, University of Oxford, 11a Mansfield Road, Oxford, OX1 3SZ, U.K.; Department of Animal & Plant Sciences, University of Sheffield, Alfred Denny Building, Western Bank, Sheffield, S10 2TN, U.K.; Department of Biology, University of Ljubljana, 1000 Ljubljana, Slovenia; Department of Ecology and Evolutionary Biology, Cornell University, Ithaca, New York, 14850, U.S.A.; Institute of Evolutionary Ecology and Conservation Genomics, University of Ulm, Ulm, 89081, Germany; Department of Biology, Case Western Reserve University, Cleveland, Ohio, U.S.A.; Department of Archaeology, University of Cambridge, Cambridge, CB2 3DZ, U.K.; College of Life and Environmental Sciences, University of Exeter, Penryn, Cornwall, TR10 9EZ, U.K.; Institute for Biodiversity and Ecosystem Dynamics, University of Amsterdam, Amsterdam, The Netherlands; Branch of Species Status Assessment Science Support, U.S. Fish and Wildlife Service, U.S.A.; School of Biological Sciences, University of Bristol, Bristol, U.K.; Institute of Biology, Martin Luther University, Halle-Wittenburg, Halle (Saale), Germany; German Centre for Integrative Biodiversity Research (iDiv) Halle-Jena-Leipzig, Leipzig, Germany; Department of Biology, Tufts University, Medford, MA, 02155, U.S.A.; School of Ocean and Earth Science, University of Southampton, Southampton, SO14 3ZH, U.K.; Department of Community Ecology, Helmholtz Centre for Environmental Research-UFZ, Halle (Saale), Germany; Department of Biology, University of Southern Denmark, Odense, Denmark; Animal Ecology and Physiology, Radboud University, 6500 GL Nijmegen, The Netherlands; NIOO-KNAW, Animal Ecology, 6700 AB Wageningen, The Netherlands; Veterinary Department, Cats Protection, National Cat Centre, Haywards Heath RH17 7TT, U.K.; Bristol Veterinary School, University of Bristol, Bristol BS40 5DU, U.K.; School of Biological Sciences and Department of Mathematics, University of Nebraska, Lincoln, NE 68588, U.S.A.; Alexander Center for Applied Population Biology, Conservation & Science Department, Lincoln Park Zoo, Chicago, Illinois, U.S.A.; School of Natural Resources, University of Nebraska-Lincoln, Lincoln, NE, 68583-0974, U.S.A.; Global Resistance Management Team, Bayer U.S. – Crop Science, Chesterfield, MO 63017, U.S.A.; Department of Biology, University of Turku, Turku, Finland; School of Life Sciences, University of Lincoln, Brayford Pool, Lincoln, LN6 7TS, U.K.; Department of Biology, University of Puerto Rico, San Juan, PR, 00925-2537, U.S.A.; Center for Applied Tropical Ecology and Conservation, University of Puerto Rico, San Juan, PR, 00925-2537, U.S.A.; Department of Biology, University of Puerto Rico-Humacao, Humacao, PR, 00791, U.S.A.; Interdisciplinary Centre on Population Dynamics, University of Southern Denmark, Denmark; Centre for Biodiversity and Conservation Science, University of Queensland, St Lucia, 4071, QLD, Australia; Evolutionary Demography Laboratory, Max Planck Institute for Demographic Research, Rostock, 18057, Germany

**Keywords:** comparative demography, matrix population models (MPM), open access, reproducibility

## Abstract

1. Stage-based demographic methods, such as matrix population models (MPMs), are powerful tools used to address a broad range of fundamental questions in ecology, evolutionary biology, and conservation science. Accordingly, MPMs now exist for over 3,000 species worldwide. These data are being digitised as an ongoing process and periodically released into two large open-access online repositories: the COMPADRE Plant Matrix Database and the COMADRE Animal Matrix Database. During the last decade, data archiving and curation of COMPADRE and COMADRE, and subsequent comparative research, have revealed pronounced variation in how MPMs are parameterized and reported.
2. Here, we summarise current issues related to the parameterisation and reporting of MPMs that arise most frequently and outline how they affect MPM construction, analysis, and interpretation. To quantify variation in how MPMs are reported, we present results from a survey identifying key aspects of MPMs that are frequently unreported in manuscripts. We then screen COMPADRE and COMADRE to quantify how often key pieces of information are omitted from manuscripts using MPMs.
3. Over 80% of surveyed researchers (n=60) state a clear benefit to adopting more standardised methodologies for reporting MPMs. Furthermore, over 85% of the 300 MPMs assessed from COMPADRE and COMADRE omitted one or more elements that are key to their accurate interpretation. Based on these insights, we identify fundamental issues that can arise from MPM construction and communication and provide suggestions to improve clarity, reproducibility, and future research utilising MPMs and their required metadata. To fortify reproducibility and empower researchers to take full advantage of their demographic data, we introduce a standardized protocol to present MPMs in publications. This standard is linked to www.compadre-db.org, so that authors wishing to archive their MPMs can do so prior to submission of publications, following examples from other open-access repositories such as DRYAD, Figshare, and Zenodo.
4. Combining and standardising MPMs parameterized from populations around the globe and across the tree of life opens up powerful research opportunities in evolutionary biology, ecology, and conservation research. However, this potential can only be fully realised by adopting standardised methods to ensure reproducibility.

## Introduction

Population ecology has come of age. The development of theories (*e.g.*, Tuljapurkar, 1982; Pfister, 1998), experimental approaches (Pradel, 1996; Crone et al., 2011) and statistical methodologies (Caswell, 2001, 2010) have resulted in the publication of demographic information for an increasingly representative sample of the world’s biodiversity (Wilmoth, Andreev, Jdanov, & Glei 2007; De Magalhᾶes & Costa, 2009; Jasilioniene et al., 2015; Salguero-Gómez et al., 2015, 2016; Levin et al., 2022; DATLife [Max Plank Institute of Demographic Research 2022; https://datlife.org/])). These data span the taxonomic tree from microbes (Jouvet, Rodríguez-Rojas, & Steiner, 2018) to macro-vertebrates (Fujiwara & Caswell, 2001; Holmes & York, 2003), and cover virtually all continents and biomes – though with important taxonomic biases (Conde et al., 2019; Römer, Dahlgren, Salguero-Gómez, Stott, & Jones, 2021). The potential of this impressive and rapidly increasing amount of information is starting to be realised. Indeed, through combining these demographic models, researchers have identified functional traits that explain variation in plant life history strategies (Adler et al., 2014; also see Bernard et al., 2022), short-term (transient) characteristics that drive the demographic dynamics of plant populations in variable environments (McDonald, Stott, Townley, & Hodgson, 2016), and ways in which life history strategies allow species to persist alongside a changing climate (Paniw, Ozgul, & Salguero-Gómez, 2018; Paniw, Maag, Cozzi, Clutton-Brock, & Ozgul, 2019; Jelbert et al., 2019; Hernández-Yáñez, Kim, & Che-Castaldo, 2022; Jackson, Le Coeur, & Jones, 2022; Le Coeur, Yoccoz, Salguero-Gómez, & Vindenes, 2022).

One of the most widely used tools for describing and analyzing species’ complex life histories is the matrix population model (MPM, hereafter). Briefly, in an MPM, individuals of a population are classified by discrete stages and/or ages (st/age hereafter) according to some biological (Caswell, 2001, p. 31) or statistical/sampling criteria (Vandermeer, 1978; Moloney, 1986; Salguero-Gómez & Plotkin, 2010). These individuals are followed in discrete time steps, typically adjusted by the generation time of the species. Indeed, time steps can vary from 12-24h as in nematode worms Caenorhabditis elegans and aphids Myzus periscae (Li, Ju, Liao, & Liao, 2014; Bruijning, Jongejans, & Turcotte, 2019), to monthly/annually periods in mammals and plants (Coulson et al., 2001; Schödelbauerová, Tremblay, & Kindlmann, 2010; Ferreira, Kajin, Cerqueira, & Vieira, 2016), all the way to 50 years in slow-growing red woods (Namkoong & Roberds, 1974). From these data, researchers estimate losses through mortality, transition probabilities among st/ages and their per-capita a/sexual contributions via reproduction (Mandujano & Escobedo-Morales, 2008; Nordstrom, Dykstra, & Wagenius, 2021; Omeyer et al., 2021).

A single MPM can be used to calculate a vast repertoire of biologically meaningful outputs. These outputs include proxies for the performance and viability of populations, such as deterministic (*λ*) or stochastic population growth rates (*λ*_S_) (Doak, Morris, Pfister, Kendall, & Bruna, 2005), quasi-extinction risk (Davis, 2022), population response to perturbations to underlying vital rates such as survival or reproduction (Caswell, 2001, p. 206), transient dynamics (Tenhumberg, Tyre, & Rebarber, 2009; Ezard et al., 2010; Stott, Townley, & Hodgson, 2011; Capdevila, Stott, Beger, & Salguero-Gómez, 2020), effective population size (Orive, 1993), and life history traits, such as rates of senescence (Baudisch et al., 2013; Caswell & Salguero-Gómez, 2013), degree of iteroparity (Salguero-Gómez et al., 2017), and age at maturity (Caswell, 2001, p. 124). This wealth of demographic inference highlights why many advances in demography and life history theory frequently utilize MPMs (Colchero et al., 2019; Franco & Silvertown, 1996; Morris & Doak, 2004; Pfister, 1998; Sæther et al., 2013; Tuljapurkar, 1989).

MPMs for plants and animals have been archived, error-checked, complemented with additional information (*e.g.*, GPS coordinates, IUCN conservation status), and released open-access in the COMPADRE Plant Matrix Database (Salguero-Gómez et al., 2015) and the COMADRE Animal Matrix Database (Salguero-Gómez et al., 2016). In the latest data release, COMPADRE v. 6.22.5 [COMADRE v. 4.21.8] contains 8,851 [3,317] MPMs from 760 [415] unique species published in 643 [395] studies. At the time of writing, a further 1,307 species are pending digitization in the COMPADRE network, at a rate of 4.5 new works containing MPMs being screened, digitized, and quality checked every week (S. Gascoigne, pers. obs.). However, one of the challenges of the digitization process – which underpins the potential of comparative demography (Buckley & Puy, 2022) – is the tremendous variation in how data are collected, presented and used to parameterize MPMs.

Data standardisation improves reproducibility and promotes data sharing across research disciplines (Reichman, Jones, & Schildhauer, 2011; Augusiak, van den Brink, & Grimm, 2014). Data standardization is therefore key for research to be replicated, validated, openly discussed, and ultimately for science to advance (Gilliland, 2008; Baker & Millerand, 2010; Reichman et al., 2011; Qin, Ball, & Greenberg, 2012; Peres-Neto, 2016; Powers & Hampton, 2019; Salguero-Gómez, Jackson, & Gascoigne, 2021). Examples of these standards include reporting sample size and variance of estimates and detailing the full list of original sources of data (Gerstner et al., 2017). In this context, standards can be used as checklist items to improve publications quality and reproducibility (Reichman et al., 2011) and to aid the peer-review process (Peres-Neto, 2016). Furthermore, meta-analyses (Gurevitch & Hedges, 1999; Hedges, Gurevitch, & Curtis, 1999; Gurevitch, Koricheva, Nakagawa, & Stewart, 2018) and phylogenetic comparative analyses (Salguero-Gómez et al., 2017; Healy, Ezard, Jones, Salguero-Gómez, & Buckley, 2019; Bernard, Compagnoni, & Salguero-Gómez, 2020), which offer valuable opportunities to examine general patterns and identify gaps in knowledge, rely on data conforming to certain standards.

MPMs are being adapted, extended, and applied beyond their original, species-specific context in comparative demography. However, not all MPMs are built and reported equally. The current presentation of MPMs in COMPADRE and COMADRE may give the false impression that all MPMs are published in a homogeneous format, despite differences in how and why the MPMs are produced (Caswell, 2001). This impression may have emerged from the amount of verification the COMADRE and COMPADRE digitisation team does behind the scenes (*e.g.*, validating model outputs, author correspondence for additional information). Whilst verification is an inevitable aspect of database curation, most of our efforts are spent communicating with authors rather than digitizing data. Our goal here is to (i) present the current standard of MPM communication in the literature, (ii) identify common issues in MPM communication and their impacts, (iii) suggest ways to support the clear communication of MPM data and metadata, (iv) highlight advantages for authors and the scientific community at large, and (v) introduce a standard method for sharing MPM data and metadata.

## MPM communication: Current state of affairs

To present the current practices in MPM data and metadata communication, with the ultimate goal to evaluate the need for standardized data and metadata reporting, we performed a survey of researchers and screened a subset of papers from COMPADRE and COMADRE.

### A survey on matrix communication

We surveyed expert population ecologists, who we identified as having published peer-reviewed papers, regarding our current ability to communicate MPM data and metadata for reproducibility purposes. Specifically, we asked how well peer-reviewed publications relay the attributes of MPMs necessary for reproducibility. Additionally, we asked if researchers thought a standardized method of matrix communication is “necessary for the coherent communication of MPMs in the literature” (the full list of 11 questions can be found in S1 in the Supplementary Online Materials). The survey was distributed using Google Forms. We identified 1,390 potential participants based on the criterion of being the lead and/or corresponding author from a publication containing at least one MPM. Over 50% of corresponding email addresses were outdated and not contacted further. Of the remaining approximately 650 researchers, that were contacted, 60 participants completed the survey. As expected, researchers report a great deal of heterogeneity in components of MPM communication (Fig. 1). The best communicated attributes according to these survey participants are trait names (*i.e.*, the phenotype by which the MPM was structured – stage/age/size classes), census duration and projection interval whilst the worst communicated attributes are life cycle graphs, formulae defining the vital rates, and population vectors (*i.e.*, number/frequency of individuals in each st/age). Importantly, 83% of survey participants agreed that the discipline needs a standardised method for MPM communication.

**Figure 1.**
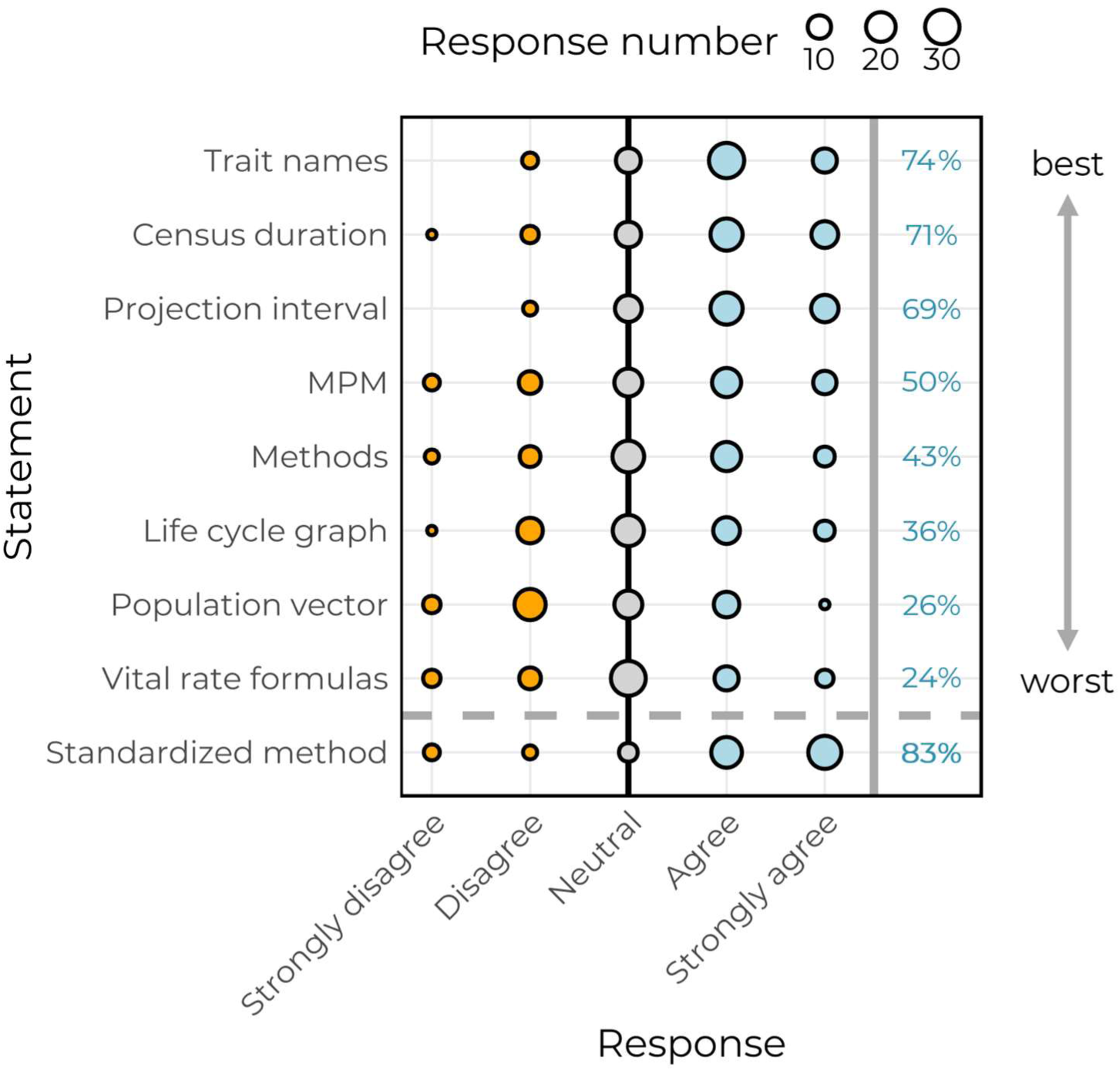
Survey results from experts in population ecology that participated (n=60). Participants ranked their confidence in the appropriate communication of components of matrix population models (MPMs) in peer-reviewed papers. Each component of MPM communication on the y-axis represents a statement shown in the survey (see S1 for the full survey). For all statements above the dotted line, participants were asked if that attribute (*e.g.*, projection interval) is sufficiently well-reported in peer-reviewed publications. The statement “standardized method” indicates participants’ response to whether the field of population ecology would benefit from a standardized method of MPM reporting. The size of the dots indicates the number of respondents with that response and are coloured (*i.e.*, orange = disagreement; grey = neutral; blue = agreement). For ease, percent agreement (*i.e.*, the percentage of participants that either agreed or strongly agreed with the statement) is shown on the right-hand side of the plot.

### A screen of papers in COMPADRE and COMADRE

To quantify how well MPM data and metadata are communicated in peer-reviewed publications, we screened 300 randomly sampled papers already digitized in COMPADRE and COMADRE (150 papers each). Across the different key attributes of MPMs that we examined, there was considerable variation in how reliably authors provided the data and metadata necessary for digitizing, archiving, and performing comparative analysis (Fig. 2). For instance, the generic location of the examined population (*i.e.*, province/city/landmark; COMPADRE: 95.1%, COMADRE: 86.2% of papers reported it), the fully-parameterized MPM (93.3%, 88.9%), and the census date (89.6%, 77.7%) were frequently explicitly stated in the papers, whilst latitude-longitude of the examined population (52.4%, 39.9%), its life-cycle diagram (44.5%, 40.1%), and population vector (*i.e.*, st/age distribution of individuals at time t) (33.2%, 32.6%) were not. Interestingly, plant studies using MPMs (COMPADRE) contain overall more explicit data and metadata than animal studies (COMADRE; Fig. 2). Furthermore, we used this information to categorise the quality of each of the examined 300 papers according to their reproducibility – from inadequate to an exhaustive inclusion of components of MPM communication (Fig. 3). The distribution of component communication across kingdoms is similar. Crucially, only 13.9% of papers in COMADRE and 15.8% of papers in COMPADRE contain all the information necessary for comparative analyses and accurate projections (Fig. 3). Thus, approximately 85% of papers require emailing authors to request undisclosed information.

**Figure 2.**
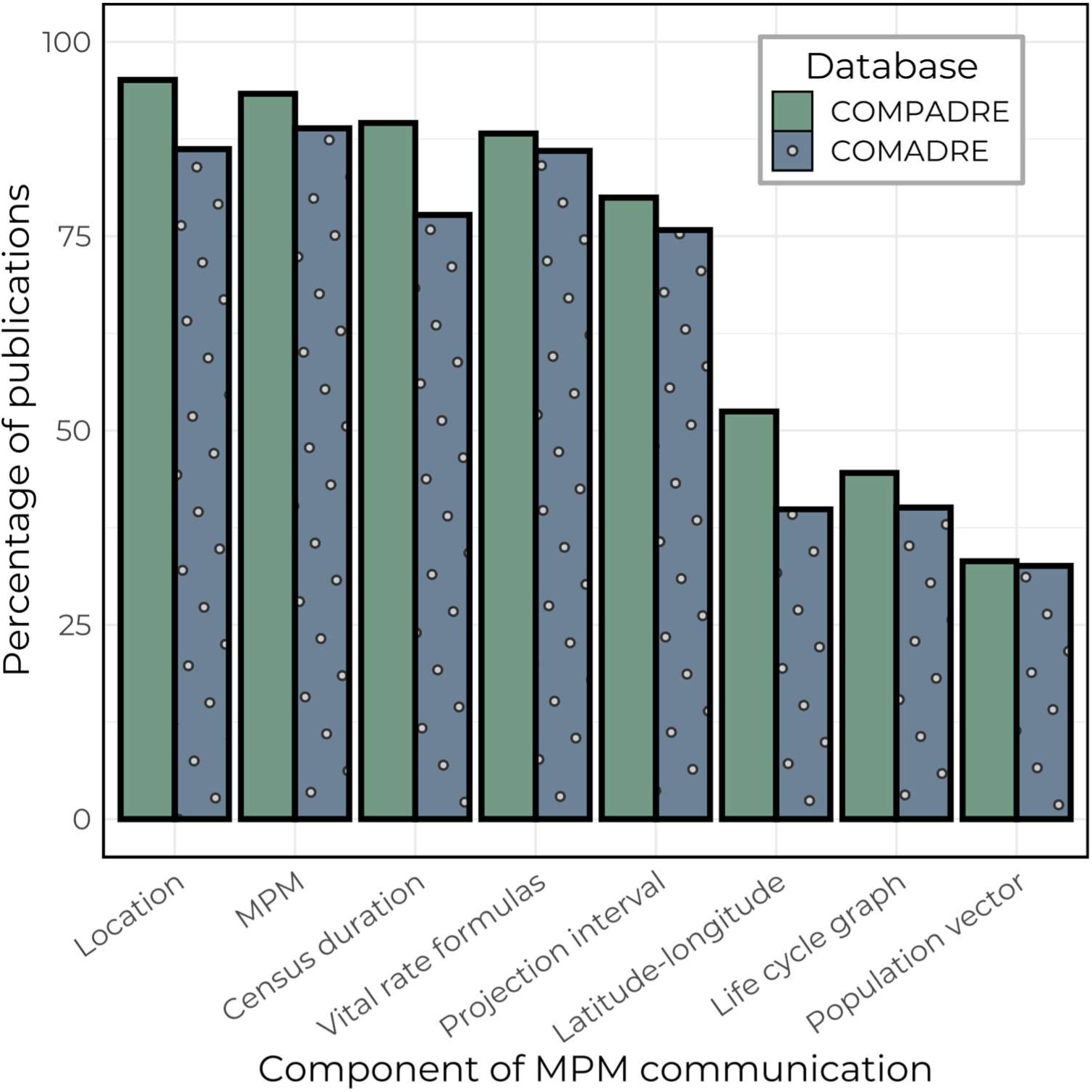
Both plant and animal MPM papers show similar heterogeneity presented components of MPM. Percentage of peer-reviewed publications in COMPADRE and COMADRE that contain a given attribute necessary for the clear communication of MPM information and its reproducibility from a random subset of 150 papers of the 643 total papers from COMPADRE and 150 out of the 395 total papers from COMADRE (300 papers total). The attributes are: Location: province/city/landmark; MPM: was the MPM explicitly reported; Census duration: start and end dates for data collection; Vital rate formulas: decomposition of matrix elements into their underlying components (*i.e.*, contributions from survival, growth, and reproduction); Projection interval: the time period between observations; Latitude-longitude: spatial coordinates; Life cycle graph: the visual representation of demographic transitions and a/sexual per-capita contributions; Population vector: st/age distribution of individuals at time t associated with reported MPMs.

**Figure 3.**
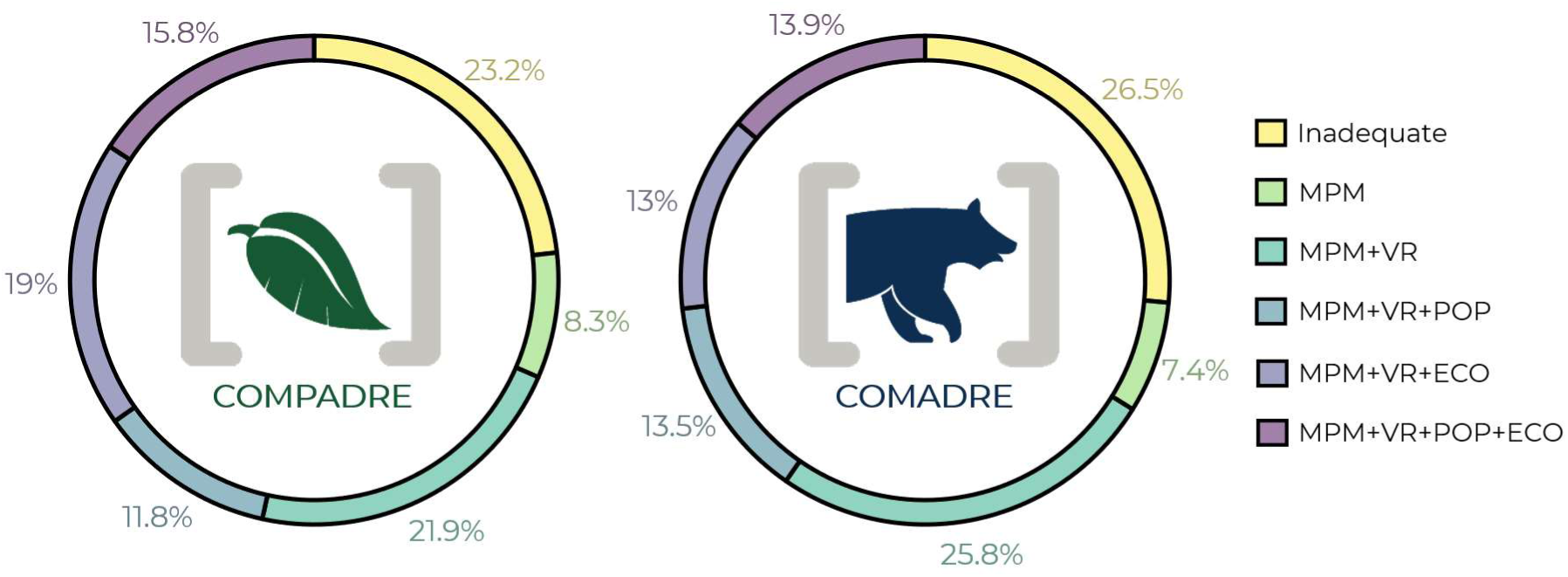
Across plant and animal MPM papers, most publications do not contain sufficient information for reproducibility. Proportion of papers in COMPADRE and COMADRE grouped by their open-access information in peer-review publications regarding matrix population model (MPM) data and metadata. Following the same scheme as in Figure 2, papers were ranked into six groups from “inadequate” to “MPM+VR+POP+ECO” (*i.e.*, fully reproducible). “Inadequate” refers to papers missing the MPM and/or projection interval (*i.e.*, an MPM specific time interval necessary for projection), without which most demographic outputs cannot be calculated. “MPM”: paper contains the MPM and projection interval but no vital rate formulas describing the matrix elements. “MPM+VR”: contains all of the information for “MPM” along with vital rate formulas for the matrix elements. “MPM+VR+POP”: contains all of the information for “MPM+VR” along with the population vector. “MPM+VR+ECO”: contains all of the information for “MPM+VR” along with latitude-longitude coordinates and census duration of the examined population. “MPM+VR+POP+ECO”: contains all of the information for “MPM+VR” along with population vector/distribution, latitude-longitude coordinates and census duration.

## Common issues in matrix construction

Here, we identify key issues in the parameterization of MPMs to illustrate the impact of methodology on demographic inference. To do so, we draw from the findings from the previous section and our experience curating COMPADRE and COMADRE. We outline the following issues for two reasons: (i) to advise demographers in how to identify them in the literature and (ii) to prevent theses issues persisting in future publications. We note that a comprehensive list was recently made available by Kendall et al. (Kendall et al., 2019 see also Che-Castaldo et al. 2020). Here, we add to these previous papers by outlining steps for researchers to avoid/mitigate these issues in their own research. A summary of these issues, from occurrence to impact, is detailed in Figure 4.

**Figure 4.**
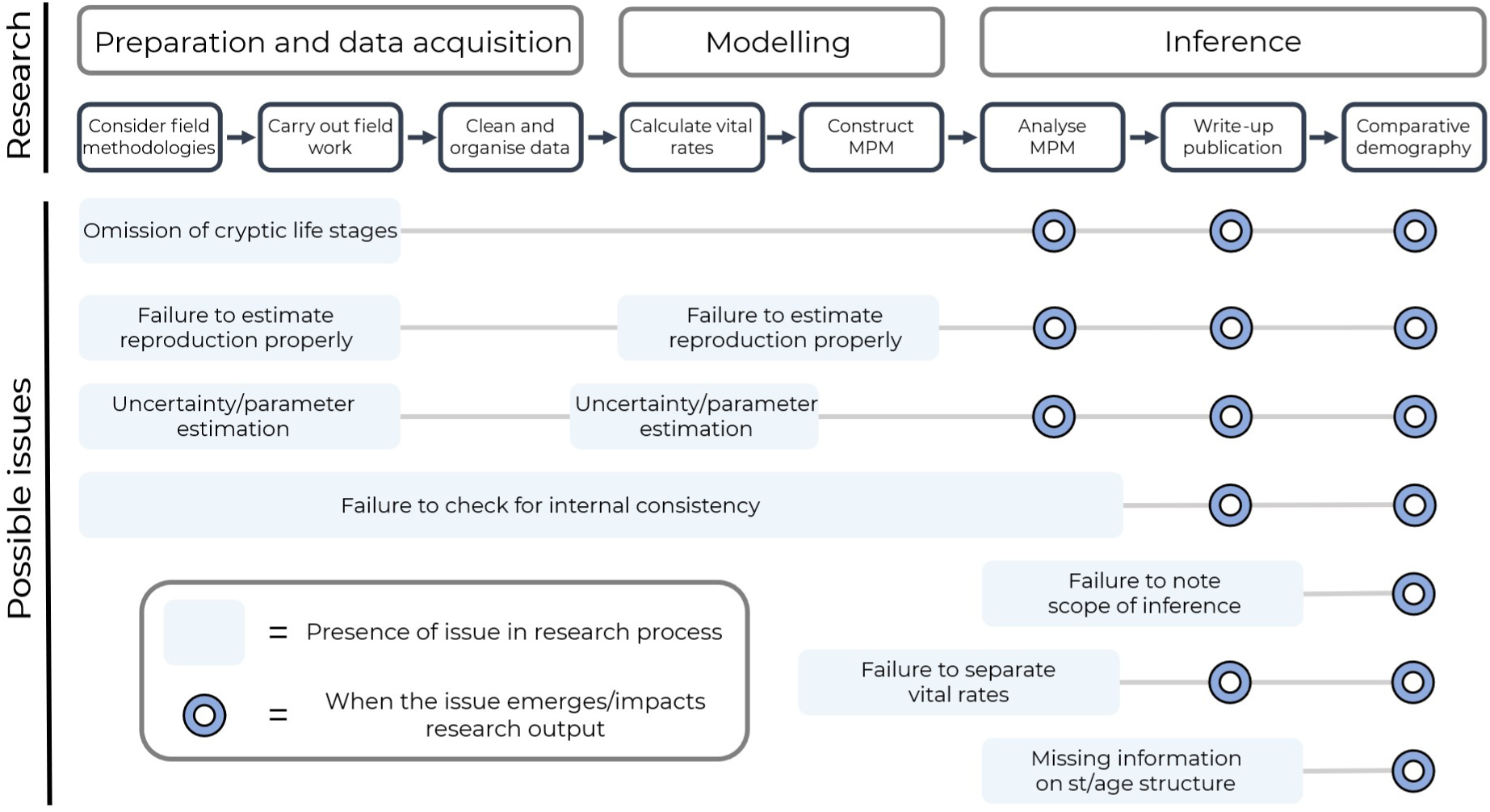
The current workflow for the production and scaling of demographic information using matrix population models (MPMs). We block the workflow into three main sections: “Preparation and data acquisition” for construction of methodologies to lab/field work, “Modelling” for the coercion of data into MPMs, and lastly “Inference” for the interpretation of demographic outputs for current and future publications. Along this workflow, we highlight when potential issues can occur, and where their consequences can emerge in demographic analyses.

### Census type, timing and frequency

MPMs are discrete-time demographic models parameterized by the tracking of individuals across censuses. Thus, the type (*e.g.*, longitudinal, cross-sectional), timing, and frequency of sampling needs to be carefully planned. These criteria are particularly important as census type directly affects matrix construction (Okuyama, 2019), and census timing and frequency can inadvertently influence demographic outputs (Emery & Gross, 2005).

Typically, an MPM comes in two forms regarding the spread of reproduction between censuses: birth-flow or birth-pulse (see Caswell, 2001, p. 22). The distinction is based on whether reproduction occurs continuously (*i.e.*, birth-flow) or in a narrow temporal window (*i.e.*, birth-pulse). Birth-pulse MPMs are further categorized into pre- vs. post-reproductive census. Although both pre- and post-reproductive censuses often lead to similar demographic inference (see Cooch, Gauthier, & Rockwell, 2003), their difference lies in when populations are censused relative to the position of the narrow reproductive window. In the former, populations are censused immediately before a reproductive window, whilst post-reproductive censuses follow on from a reproductive window. A pre-reproductive census requires the inclusion of offspring survival in reproductive matrix elements, whilst a post-reproductive census requires the inclusion of parent survival in reproductive matrix elements. We often encounter mistakes in the accommodation of offspring or parent survival in reproductive matrix elements (see also Kendall et al., 2019). A key step in matrix construction that can prevent the incorrect accommodation of survival is drawing the life cycle graph (as per Ebert, 1999, p. 61) with respect to census timing (demonstrated in Ellner, Childs, & Rees, 2016, p. 13), as well as explicitly detailing the census type used to parameterize the MPM. However, sometimes drawing the life cycle graph may be unfeasible or uninformative. For example, the graph for an age classified model with 100 age classes is too large to draw and too redundant to be useful. Models with many stages and highly connected transitions are not feasible to draw the life cycle graph (*e.g.*, the graph for Calathea ovandensis in Neubert and Caswell (2000)). But even in complex situations (*e.g.*, the series of seasonal graphs for the emperor penguin in Jenouvrier et al. (2010)) the graph may be helpful in organizing the structure of the model.

Census timing and frequency affects model construction, making a constructed MPM impractical for demographic inference if the life history of the examined organism is not considered. Consider a researcher comparing the demographic processes of fruit flies and fruit trees. The researcher first notices that there are four discrete stages to the fruit flies’ life history: three juvenile stages encompassing the development from egg to instar to pupae, and one adult stage where individuals disperse and reproduce. Since development from egg to adult takes ~10 days in this species, the researcher decides to perform the census every 10 days for both the fruit fly and the fruit trees over a three-month period. However, because neither mortality nor reproduction occur across such a short census in the fruit tree population, the resulting fruit tree MPM, when projected forward, will persist forever, neither increasing nor declining. This same issue would occur the other way around. If five-year intervals were deemed sufficient for the fruit trees, then individually measured fruit flies would never survive across time steps.

A solution to this problem exists, using periodic matrix models to include periods much shorter or longer than other periods. For example, Hunter and Caswell (2005) analyzed the Sooty Shearwater (Puffinus griseus) including two harvesting periods of several weeks in duration and then an annual interval for the species, with a lifespan of decades. Smith et al. (2005) and Shyu et al. (2013) used periodic seasonal models to accommodate life cycles in which some stages are only present for part of the annual cycle. The approach (Caswell 2001, Section 13.1) is powerful and general.

### Unrealistic stage-specific survival

Issues in parameterising stage-specific survival, whilst easy to diagnose, can result in an array of unnatural life histories. Transition and survival probabilities are bounded between 0 (*i.e.*, the event never happens) and 1 (*i.e.*, always occurs). As such, the stage-specific survival of an MPM, the summed non-reproductive elements in a given column of the MPM ***A*** must not exceed 1. When it does, individuals in that stage have an unrealistic average chance of surviving >100%, resulting in an incorrect representation of the organism’s life history. Stage-specific survival values >1 typically arise due to rounding errors, typos, inclusion of unstated a/sexual reproductive events. As such, it is generally advised to omit these MPMs in comparative analysis (Jones et al., 2014). Unstated a/sexual reproductive events occur when a given element *a_i,j_* in the MPM ***A*** contains both survival-dependent processes, such as growth/shrinkage, but also fertility, and these have not been reported separately. Ideally, authors would carefully identify whether various vital rates are being confounded with survival-dependent demographic processes in each MPM element. For the comparative demographer using COMPADRE and COMADRE, we recommend either avoiding MPM models where stage-specific survival >1 or altering the model so that the stage-specific survival is fixed to a maximum of 1 (*e.g.*, Buckley et al., 2010) However, prior to altering an MPM in this way, we suggest a sensitivity analysis of the output of interest to assess the effect of this alteration.

In many published MPMs, some life stages have a survival probability of 1 or incomplete life cycle, likely the result of small sample size or rare event along the life history of the species. Perfect survival (*i.e.* mortality = 0) is unlikely to be accurate, and may need to be estimated or imputed (Johnson et al., 2018).

A reproducible approach to infer realistic survival and transition values was recently proposed by Tremblay et al. (2021), using a Bayesian approach to estimate parameter values using priors in addition to the observed data to obtain posterior MPMs. An advantage of this approach is that the confidence intervals of the parameters (*i.e.*, stasis, transition, survival) have a beta distribution. This advantage of using a Bayesian inferred multinomial Dirichlet distribution for estimating the mean values is that the researchers can infer variance and skew of the posterior distributions to further inform for MPM construction and demographic inference.

### Incorrectly parameterizing fertility

Fertility often presents a challenge to constructing accurate MPMs. This challenge is partly due to the ambiguity of the term ‘“fertility”. The issue arises when the per-capita contributions of reproductive adults to new recruits (*e.g.*, eggs, neonates, seeds, etc.) do not represent the links over the full projection interval of the study. Remember that the entry *a_i,j_* in an MPM is the (expected) number of stage *i* individuals at *t* + 1 per stage *j* individual at time *t*. If stage *i* is some kind of “newborn” individual, then *a_i,j_* must include all the processes between time *t* and time *t* + 1 (Caswell, 2001, pg. 61). Reproductive output, in turn, is a composite demographic process of the number of offspring produced in a reproductive event and the relevant survival that will penalise how many new offspring will actually make it to the next observation. Failure to accommodate this vital rate decomposition can result in the introduction of a one-timestep lag into the organism’s life cycle, as newly created offspring spend a projection interval “in limbo” before their onward transitions. The best-known example is in the classic model of teasel (Dipsacus sylvestris) by Werner and Caswell (1977), in which flowering plants at time *t* were described as producing seeds at time *t* + 1, which only germinated to seedlings at time *t* + 2. The issue was discussed and corrected in Caswell (2001). Furthermore, this issue has been reported, for instance, in reproductive structures such as seeds that do not actually undergo a permanent seed bank. An MPM with this issue will typically (Kendall et al., 2019), but not always (Nguyen, Buckley, Salguero-Gómez, & Wardle, 2019), underestimate the asymptotic population growth rate, *λ*. Naturally the challenge will then be in estimating the relative importance of the seed bank and the lifespan on non-germinated seeds. The effect of incorrectly parameterizing fertility on *λ* is greatest in cases of extreme growth, such as invasive species, or extreme decline, such as critically endangered species (Rueda-Cediel, Anderson, Regan, & Regan, 2018). Furthermore, this issue can also cause overestimation of the transient envelope (see Ezard et al., 2010). Thus, we recommend reporting the fertility vital rate formulas with the associated MPMs and clearly identifying the values of these underlying vital rates (as in Box 1).

### Indirectly calculating vital rates

Estimating vital rates often involves combinations of direct and indirect measurements. Direct measurement empirically derives vital rates from individual-based data where individuals are censused multiple times, as in cohort life table studies, mark-recapture methods and many quadrat studies of marked plants (*e.g.*, Hernández, van Daalen, Caswell, Neubert, & Gribble, 2020). However, vital rates can be hard to observe in species with high offspring production, complex phenology, and/or small population sizes (Beissinger & Westphal, 1998). Consequently, recruitment estimates are often supplemented into MPMs from controlled conditions; examples include the laboratory (Jouvet et al., 2018), greenhouse (Gontijo & Carvalho, 2020), zoo (Clubb et al., 2009) and botanic garden (Jiménez-Valdés, Godínez-Alvarez, Caballero, & Lira, 2010). Since some MPM methods require a full life cycle to obtain key metrics (*e.g.*, *λ*: Caswell, 2001, p. 106; transient metrics: Stott et al., 2011), external study sites or literature sources are often used to parameterize components of the MPM to “close the loop” in incomplete life cycles (Omeyer et al., 2021). However, captive populations may not represent wild population dynamics (Clubb & Mason, 2003; Johnsson, Brockmark, & Näslund, 2014), particularly in regards to survival (Jule, Leaver, & Lea, 2008; Che-Castaldo et al., 2021) or reproduction (Clubb et al., 2009).

Another method to indirectly estimate vital rates involves using ex-situ methods to obtain upper and lower bounds on recruitment (or other vital rates) and explore the parameter space within those bounds. The approach was introduced by Caswell et al. (1998) in a study of the effects of bycatch mortality on the harbour porpoise. Age-specific survival and fertility schedules were selected from other species with similar life cycles, re-scaled to match the longevity of the harbour porpoise, and used to produce uncertainty distributions for population growth and the effects of the measured bycatch. Reporting the distribution and associated parameters provide a measure of uncertainty from which to inform the construction of an MPM (Lubben, Tenhumberg, Tyre, & Rebarber, 2008; Tenhumberg, Louda, Eckberg, & Takahashi, 2008). Furthermore, the use of hierarchical models to estimate missing values and borrowing strength from other populations or species may improve parameter estimation (Tremblay et al., 2009a, 2009b; Tremblay & McCarthy, 2014; Che-Castaldo, Che-Castaldo, & Neel, 2018; James, Salguero-Gómez, Jones, Childs, & Beckerman, 2021; Davis, 2022).

### Population vector

An estimate of the structure of the population, classified by age or stage, is a useful piece of information when available, but it will only sometimes be available. It can be used as a starting point for projections of short-term, and long-term population viability (Hunter et al., 2010; White et al., 2013; Jackson, Childs, Mar, Htut, & Lummaa, 2019; Werner & Peacock, 2019). Reported population stage or age vectors often result from two key components: the actual population structure at the census in time *t*, and the methodological choices. This second component is critical to accurately represent the studied population. Across the development and curation of COMPADRE and COMADRE we have noticed two sources of bias that affect population vectors: observer bias and sampling bias. Observer bias arises when researchers identify certain st/ages with a higher rate of detection over more cryptic stages (*e.g.*, seed banks). A less discussed bias for constructing population vectors, sampling bias, is the researcher’s effort to measure more individuals from a certain st/age class over other classes. This latter point is particularly common when st/age classes have survival probabilities close to its limits (*i.e.*, 0 and 1). For instance, in tree demography, there are typically only a few very large individuals per area examined. Thus, oftentimes researchers supplement the sample size of this category by sampling outside of the predefined area (Jones & Hubbell, 2006).

Communicating population st/age vectors is critical for an accurate projection of populations that are not at their stable st/age distribution. However, this information is regrettably rarely included in publications (as shown in Figs.1,2). Even when these data are included in publications, population vectors are often vaguely communicated to the reader. Papers that report population vectors often do not identify whether the vector represents a sample size for vital rate calculation or a representation of population structure (D. Hodgson pers. obs.). By detailing the initial conditions of the population, further insights can be drawn in the form of population forecasts (Dietze, 2017), transient dynamics (*i.e.*, damping ratio and other resilience metrics (Capdevila et al., 2020)) as well as how far the population is from the stable st/age distribution (*i.e.*, the right eigenvector (***w***) corresponding to the first eigenvalue of the ***A***-matrix (Caswell, 2001, p. 106)). These analyses allow for insights that go beyond simple projection towards complex demographic inferences.

We note that many of the concerns involving population vectors (*i.e.*, abundance and stage distribution) stem from a basic misunderstanding of the difference between estimating rates and the estimation of numbers. MPMs require the estimation of rates, not numbers.

Many types of estimation do not provide any information on numbers and structure. Cohort life tables, that follow a cohort of individuals as they age, are blind to the structure of the population in which the cohort develops. Indeed, there may be no such population (*e.g.*, Hernández et al. (2020) and the entire history of laboratory cohort-based demography going back to Pearl in the 1920s (Pearl, 1925; Pearl, Miner, & Parker, 1927)). Mark-recapture estimation of rates from longitudinal data draws all its inference from the marked individuals and makes no inferences about the number and structure of the unmarked. The literature on mark-recapture methods for estimating rates recognizes that estimating population numbers is thus much more difficult than estimating rates (Lebreton, Burnham, Clobert, & Anderson, 1992) and requires different methods.

Since the estimation of MPMs is based on following marked individuals, the sampling design may follow arbitrary numbers of individuals in each stage, which thus provide no information on population structure. This is “biased” only if thought of as an attempt to estimate numbers and structure, but that is not what it tries to do.

We also recognize that some MPMs are constructed from measurements on only one portion of the life cycle, with many stages unobservable. This is typical for studies of seabirds (albatrosses, petrels, penguins, fulmars) in which the stages between fledgling and adults returning to the breeding colony are unobservable (Jenouvrier, Aubry, Barbraud, Weimerskirch, & Caswell, 2018; Jenouvrier, Desprez, et al., 2018). Powerful statistical methods have been developed to estimate the rates applying to these stages, but they provide no information on population size or structure.

Yet again, we also recognize that some MPMs are derived from long time-series of measurements (Coulson et al., 2001; Jackson et al., 2019; Paniw et al., 2019); a welcome development that is becoming more frequent. There is obviously no single vector describing such a population.

The interest in some kind of population vector is partly motivated by the belief that such a vector is needed as an initial condition for projection. But projections are always conditional and are not necessarily tied to “the” population structure. And if projection from an actual structure is desired, that initial condition may be more appropriately measured in a separate census, rather than extracted from the measurements of rates that inform the MPM.

### Omitting cryptic life stages

The identification and estimation of vital rates in cryptic stages poses a challenge in population ecology. Cryptic stages represent points along an organism’s life cycle that are somewhat hidden from or overlooked by population ecologists when building population models (Doak, Thomson, & Jules, 2002). A life stage could be cryptic because it is logistically challenging to observe or observable but indistinguishable from a similar seeming class (Nguyen et al., 2019). In plants, cryptic stages can emerge from seed banks for plants, such as orchids, where the seeds are too small to be identified in the field (Paniw, Quintana-Ascencio, Ojeda, & Salguero-Gómez, 2017; Nguyen et al., 2019) or some herbaceous perennials (*e.g.*, Astragulus scaphoides) where prolonged periods of vegetative dormancy can allow individuals to stay underground for one or more growing seasons (Gremer, Sala, & Crone, 2010; Gremer & Sala, 2013). Additionally, animals can exhibit cryptic stages by undergoing diapause, delays in development due to adverse environment conditions (Aedes albopictus: Jia et al., 2016). Pelagic seabirds (albatrosses, petrels, penguins) often spend pre-reproductive stages, sometimes of many years durations, at sea, completely cryptic until returning to the breeding colony as adults. Sophisticated multistate mark-recapture methods can provide estimates of parameters for these parts of the life cycle (*e.g.*, Jenouvrier et al. (2018), using the multi-event algorithm of Choquet et al. (2009)). Omitting a cryptic life stage can reduce the biological realism of an MPM (Nguyen et al., 2019) and alter the number of stages in the MPM, which can further impact demographic outputs (Vandermeer, 1978; Tenhumberg et al., 2009; Tremblay et al., 2009a, 2009b; Salguero-Gómez & Plotkin, 2010). In some cases, cryptic life stages will only be identified via a multidisciplinary approach including field and laboratory methods, coupled with Bayesian frameworks to integrate data and prior knowledge (*e.g.*, Paniw, Quintana-Ascencio, Ojeda, & Salguero-Gómez, 2017).

### One-sex *vs.* two-sex models

Much of demography focuses on females, under the assumptions that fertility is determined by females without limitation by males (see Caswell, 2001, p. 568). Such models may include males (*e.g.*, Hunter et al. 2010), but if reproduction is determined by female rates (*i.e.*, are female-dominant), males represent a set of stages that do not contribute to population growth. Most existing animal MPMs are female-based and female-dominant (Salguero-Gómez et al., 2016) because, given sampling biases towards mammals and birds (Conde et al., 2019) it is oftentimes not feasible, or necessary for the research question, to track male reproductive interactions (Archer, Paniw, Vega-Trejo, & Sepil, 2022). These studies typically assume a 1:1 sex ratio, sex-congruent vital rates and that reproduction is not male-limited (Tsai, Liu, Punt, & Sun, 2015; Compagnoni, Steigman, & Miller, 2017; Miller & Compagnoni, 2022). Whilst one-sex models are common in animal MPMs (currently 77% in COMADRE v. 4.21.8), care must be taken to not make assumptions about sex-ratio dependent dynamics within these systems (Archer et al., 2022) Indeed, these assumptions may not be met when any of the following are true: there is a bias in sex ratio (Archer et al., 2022), there is reproductive skew (Sky et al., 2022), or a high sensitivity of population dynamics in response to mating choice (Veran & Beissinger, 2009). Two-sex models that do not assume dominance by one sex are nonlinear and require specification of a mating function that describes fertility as a function of male and female abundance (Caswell 2001). Defining such mating functions is generally difficult or impossible, except in the particularly easy case of strict monogamy (Jenouvrier et al., 2010).

Reporting sex ratios can greatly expand the scope of a study (Shyu & Caswell, 2016a, 2016b); for example evaluating the impact of sex ratio and the Allee effect (Boukal & Berec, 2002). Unfortunately, this reporting is rarely done in work archived in COMPADRE and COMADRE. Moreover, if there are differences in vital rate values between sexes, such as survival, growth, and/or reproductive output, a one-sex MPM may not be representative of the population (Caswell, 2001, p. 568; Archer et al., 2022). In plants, reporting two-sex dynamics is even more rare (0.2% in COMPADRE v. 6.22.5). However, this low percentage likely reflects the rarity of dioecy or other mating systems with two or more sexes in plants (Bawa, 1980; Käfer, Marais, & Pannell, 2017) and the commonness of polygamous mating systems which makes male-limited reproduction rare (see Compagnoni et al., 2017; Miller & Compagnoni, 2022).

### Irreducibility and ergodicity

The property of irreducibility has implications for the eigenvalue spectrum of a matrix. These implications are well known in the literature on MPMs (Caswell, 2001). An irreducible matrix is one in which the life cycle graph is completely connected; *i.e.*, there exists a directed path from any stage to any other stage. It is sometimes asserted that reducible MPMs are somehow invalid; they are not. There are (at least) four situations in which reducible matrices naturally occur.

1. Life cycles with post-reproductive stages. The post-reproductive stages can make no contribution to the potentially reproductive stages (*e.g.*, MPMs for humans, orcas).
2. Two-sex models with dominance by one sex (usually females, but could be male). In a female-dominant model, all reproduction is credited to females. Males are produced by females, but make no contribution to the female part of the life cycle (*e.g.*, Hunter et al., (2010) for polar bears).
3. Spatial models in which dispersal is one-directional, as in river systems or oceanic currents.
4. Age x stage-classified models (Caswell, 2009; Caswell & Salguero-Gómez, 2013). In these models, reproduction produces (by definition) individuals in age class 1, but the model includes all combinations of age and stage, including impossible combinations of age class 1 and stages that do not exist at age 1.

Reducibility may or may not be easy to spot from the life cycle graph, but it can be tested numerically. The matrix ***A*** is irreducible if and only if the matrix (***I*** + ***A***)*^S^*^-1^ is positive (Caswell 2001). Irreducibility is a sufficient condition for ergodicity, guaranteeing that the population will converge to the same stable structure regardless of the initial condition. A reducible matrix may not have this property; clearly, for example, a population started with only post-reproductive individuals will not converge to the same structure as one started with some pre-reproductive individuals. With regard to ergodicity, it is also known that an MPM is ergodic if and only if all entries of its dominant left eigenvector (***v***) are positive (Stott, Townley, Carslake, & Hodgson, 2010). In short, despite appropriate model structure and correct parameterisation, demographic data may lead to reducible and/or non-ergodic matrices.

An MPM may be a correct representation of the transitions and fertilities observed in a population over the study period; however, taking a given MPM out of context (*e.g.*, not accounting for seasonality or rare events observed during the census period) risks misrepresenting the general life history of that population/species and incorrectly modelling how much different parts of the life cycle contribute to population dynamics. If researchers find this issue with their constructed MPMs, one method to fix these issues is by reducing matrix dimensionality (*i.e.*, the number of rows and columns in the MPM, and thus the number of stages in the life cycle). Thankfully, the methods for dimensionality reduction have been well described and offer an extra tool for researchers facing difficulties in irreducibility and ergodicity (Ramula & Lehtilä, 2005; Salguero-Gómez & Plotkin, 2010). Despite occasional statements that reducibility somehow renders demographic analyses invalid, in fact, interpretation of results of reducible MPMs just requires some careful thought. These approaches represent tools in the demographer’s toolbox to identify and mitigate the effects of reducibility and non-ergodicity on demographic inference.

## Full representation of an MPM in publications

In this section, we justify the need for clear presentation of MPMs in the scientific literature, suggest where to archive MPMs open-access, and discuss how these two actions benefit the original authors, publishing journal, readers, comparative demographers, and the discipline at large.

### Partitioning demographic processes

It is important to define what each matrix element in an MPM represents. Various demographic processes can overlap into the same matrix element in an MPM, particularly in species with a fast and/or plastic lifecycle relative to the MPM projection interval. For example, the value in an MPM that represents the link between large individuals at time *t* and smaller individuals at time *t* + 1 might correspond to sexual reproduction, clonal reproduction, fission, retrogression, or a composite of multiple processes (*e.g.*, Hoffmann, 1999; Weppler, Stoll, & Stöcklin, 2006). The mathematical derivations of key life history traits (*e.g.*, generation time, mean life expectancy, rate of senescence, degree of iteroparity) require that these processes be clearly separated (Jones et al., 2022). This is critical for the family of analyses based on Markov chains; the matrix ***U*** defines the transient state transitions in an absorbing Markov chain (Caswell, 2011, 2013). By reporting the underlying demographic rate structure in a life cycle diagram and its consequent full matrix population model ***A***, one can separate matrices into survival-dependent processes (*e.g.*, progression/growth, retrogression/shrinkage, fission, fusion, stasis) in the submatrix ***U***, sexual reproduction in the submatrix ***F*** and clonal reproduction in the submatrix ***C*** (Fig. 5). Importantly, both ***F*** and ***C*** submatrix elements must incorporate survival according to census type (*i.e.*, pre-/post-reproductive census).

**Figure 5.**
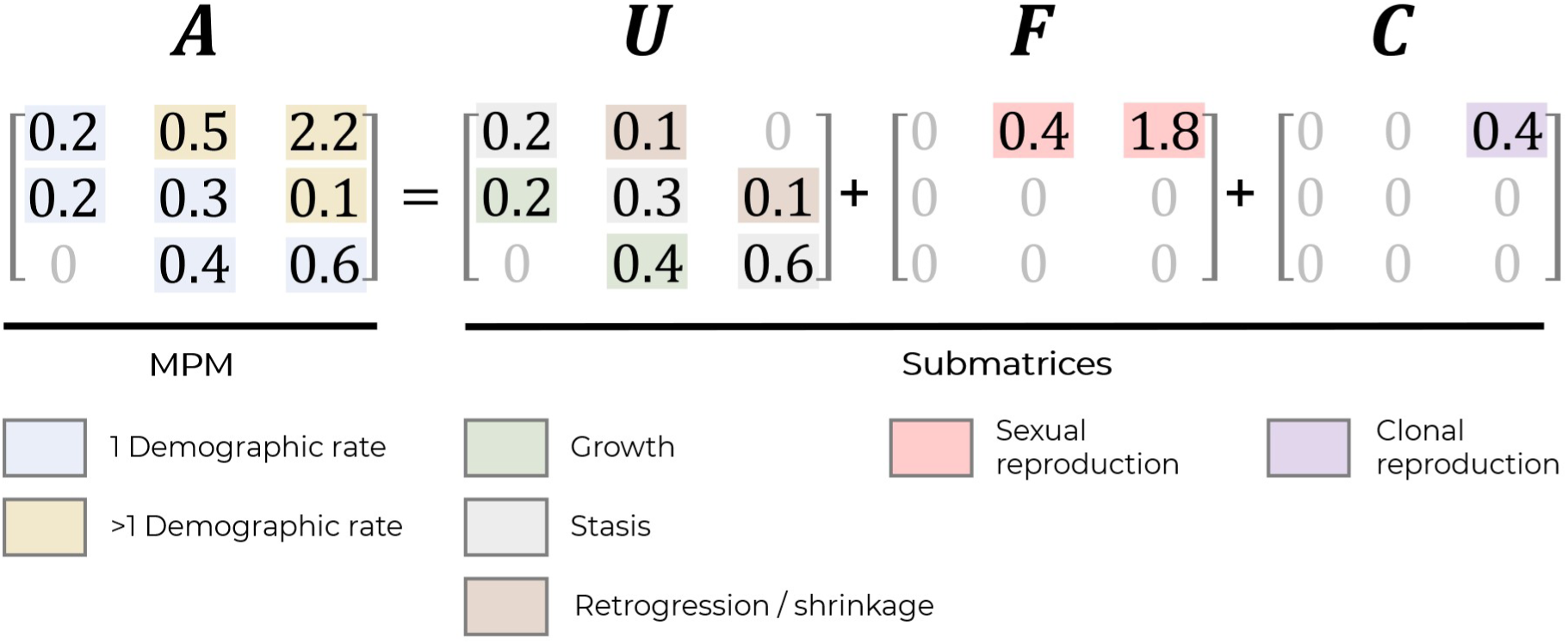
Decomposing an MPM into its submatrices allows for the isolation for otherwise masked vital rates. Matrix ***A*** represents the MPM. Since individual transitions can be represented by multiple demographic rates (*e.g.*, retrogression, sexual reproduction and clonal reproduction), decomposing ***A*** into its ***U***, ***F*** and ***C*** submatrices allows for targeted demographic inference about what demographic transitions are driving the dynamics of the population.

Reporting the matrix ***A*** as well as the submatrices ***U***, ***F***, and ***C*** lends two key benefits: (1) explicitly indicates how the values in ***A*** are generated from underlying vital rates; and (2) the submatrices can be used to calculate a vast plethora of demographic measures that cannot be calculated from ***A*** alone, such as longevity (mean and variance), occupancy times (means and variances), lifetime reproductive output (means and variances), net reproductive rate, generation time and entropy (Keyfitz entropy (Keyfitz, 1968) and Demetrius’ entropy (Demetrius, 1992)) just to name a few.

### Attribution of secondary data sources

Secondary data sources are critical for reproducibility. These data sources provide information and support for methodologies used in MPM construction. In some cases, MPMs simply use secondary data to complete the life cycle, whereas others are constructed purely from secondary sources (see Table 1). Secondary sources include previous studies, data from databases, simulations, indirect observations and theoretical estimates. The use of secondary data sources may mean that the final MPM does not accurately represent vital rate trade-offs, and so should be recognised in the methods section of the publication or its supplementary online materials. Sufficient communication of secondary sources includes both the source of the data as well as the rationale for their inclusion. For example, Omeyer et al. (2021) presents a table of data sources used in construction of the MPM. If these secondary sources are not recognized in tandem with the MPM, the inferred demographic processes, however realistic they may be, may not pass peer review nor uptake by the scientific community. In turn, clear communication of these secondary sources is highly recommended.

**Table 1.**
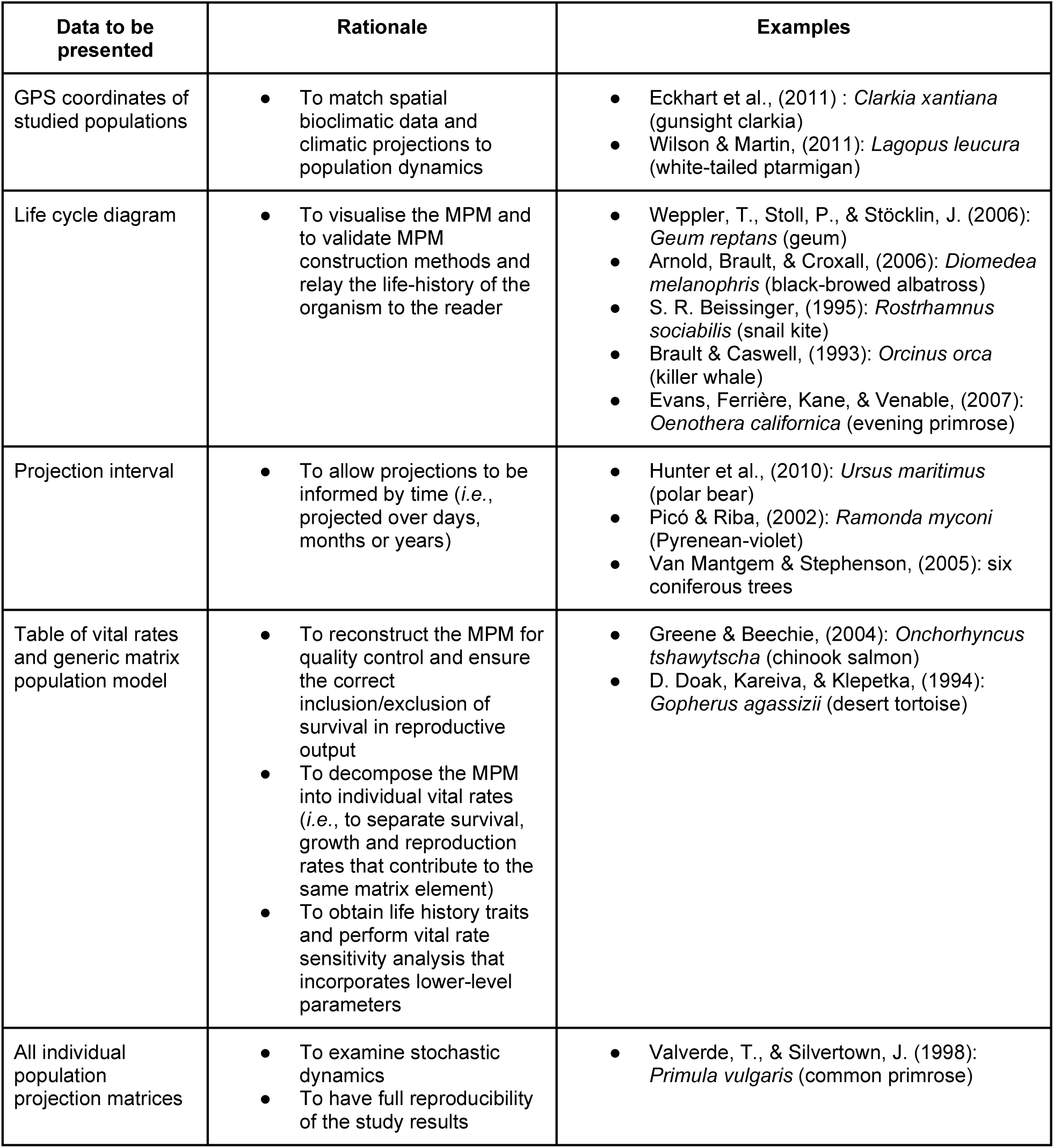
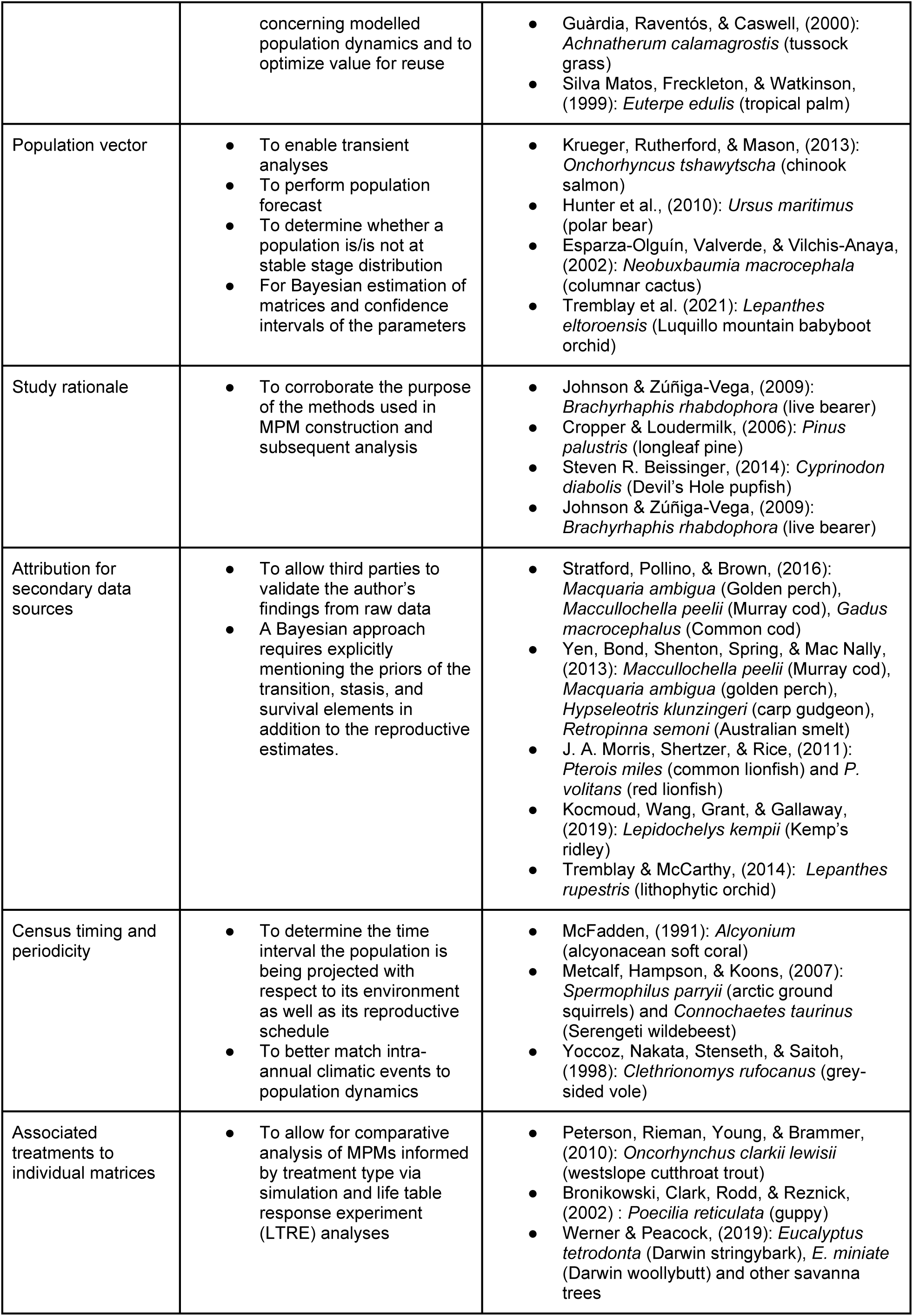
Corresponding data to include when publishing MPMs along with a rationale for inclusion and examples of good practice.

### Geographical coordinates

It can be useful to have GPS coordinates for the study populations if this is relevant and feasible. It is not always possible. Studies of wide-ranging populations (*e.g.*, whales (Brault & Caswell, 1993; Fujiwara & Caswell, 2001), polar bears (Hunter et al., 2010)) use data collected over thousands of square kilometres. Albatrosses breed in a well-defined colony, but the population is spread out literally over large portions of the globe (Laysan albatrosses nesting on Midway Island in Hawaii forage in Alaska). The environmental factors relevant to the demography of these species is not defined by the location of the breeding colony.

### Archival of information in COMPADRE and COMADRE

We propose the COMPADRE and COMADRE matrix databases provide the most appropriate way of archiving and accessing MPMs. Whilst we recognise that there are other ecological database repositories (*e.g.*, dryad: https://datadryad.org/; figshare: https://figshare.com; zenodo: https://zenodo.org), the open access COMPADRE and COMADRE databases (https://compadre-db.org/Contribute) provide a dedicated data archival platform, specifically for MPMs, allowing direct contributions from researchers as well as digitization of published MPMs by our data validation teams.

The web-based data entry portal provides a structured data curation process (*i.e.*, from screening, to standardisation, to validation) that can accommodate MPMs of different dimensions and for diverse life histories. On entry, MPMs are complemented with relevant biogeographic variables and details on census methodology in COMPADRE and COMADRE. Details of the original publication, including DOI and citation functionality (see https://compadre-db.org/Education/article/obtaining-references), are stored alongside each MPM to ensure that their contribution towards any future publication is recognised. All data are archived long-term through Oxford Open Access and Bodleian Library support.

Other recent enhancements to COMPADRE and COMADRE will further aid the research community. Previously, the databases were only accessible via download of an R-object file which contained all matrices in that version of the database. The database is now accessible via a queryable website (https://compadre-db.org/QueryDatabase) that allows users to find and download individual matrices. We also strive to empower researchers and educators with teaching materials (https://compadre-db.org/Education) and the production of new R-packages (Jones et al., 2022) for ease and scalability of MPM-related research. Along with these materials, all details of the database structure and workflow are open-access (https://jonesor.github.io/CompadreGuides/user-guide.html).

These improvements to the databases and their interface structure have been directly targeted to equip demographers with more tools to conduct research with and train students on MPMs along with increasing database transparency to ensure best research practices.

## A standard protocol for reporting MPMs

Here, we introduce a proposed checklist for how to report an MPM in publications (Box 1). We recommend using the checklist when designing data collection as well as when writing up the MPM for publication. We recommend using this template as Supplementary Material for published MPMs as it allows for the clear communication of model construction in addition to ease in integrating published MPMs into the COMPADRE and COMADRE databases.

### Box 1: Example presentation of a hypothetical three-stage plant matrix population model (MPM) Using a clear and explicit presentation of data applicable to most M PM construction techniques.

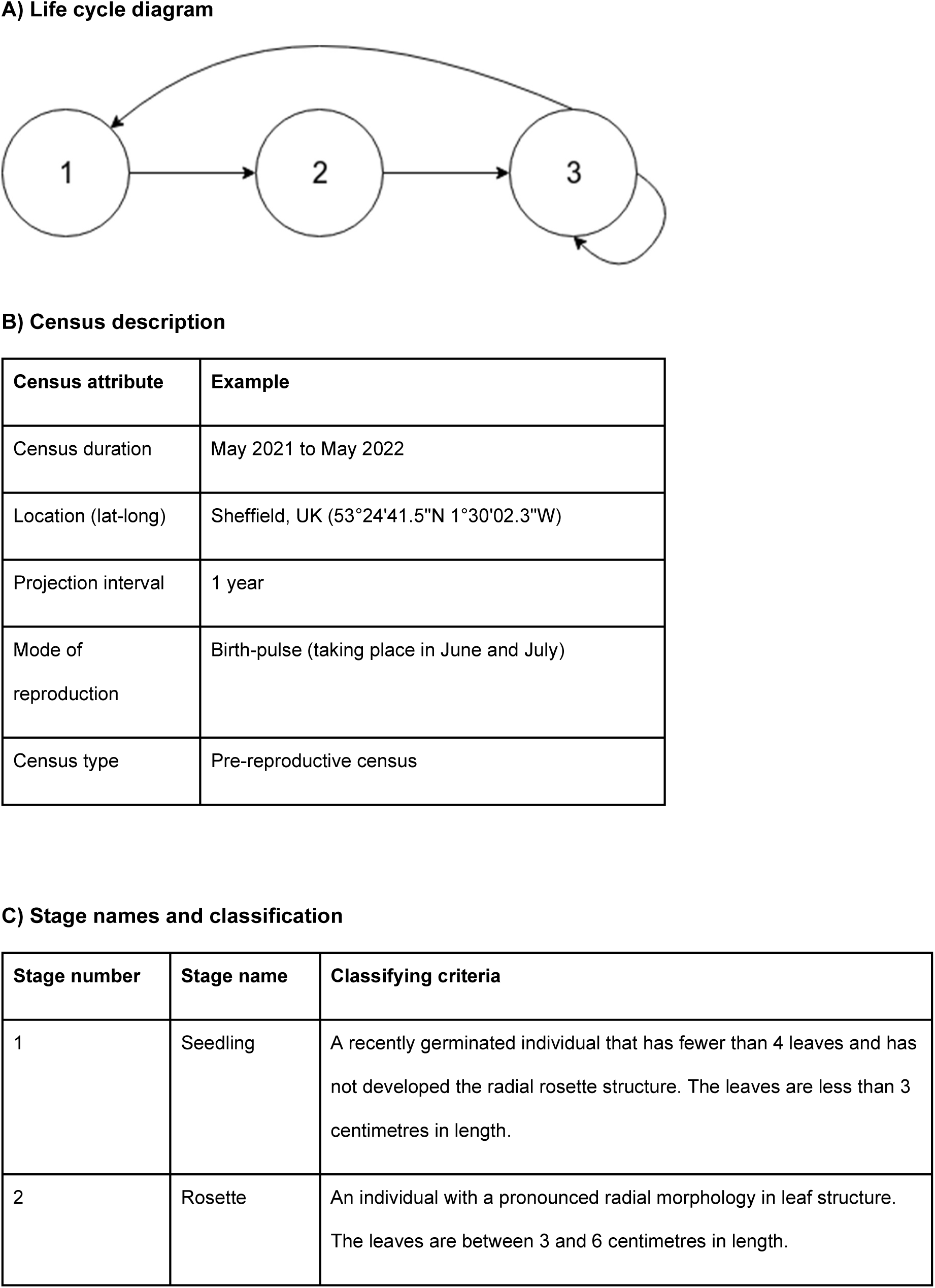
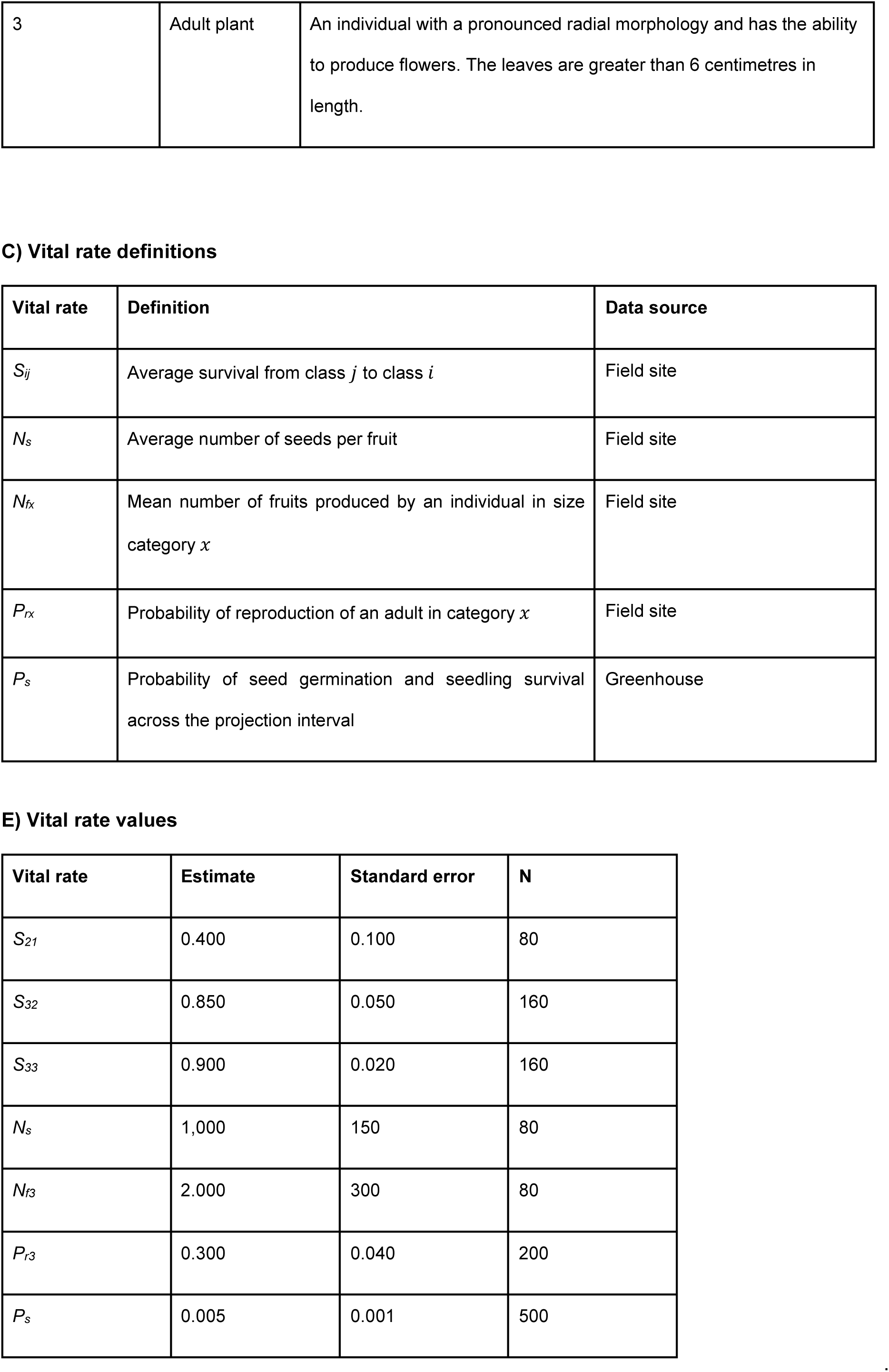
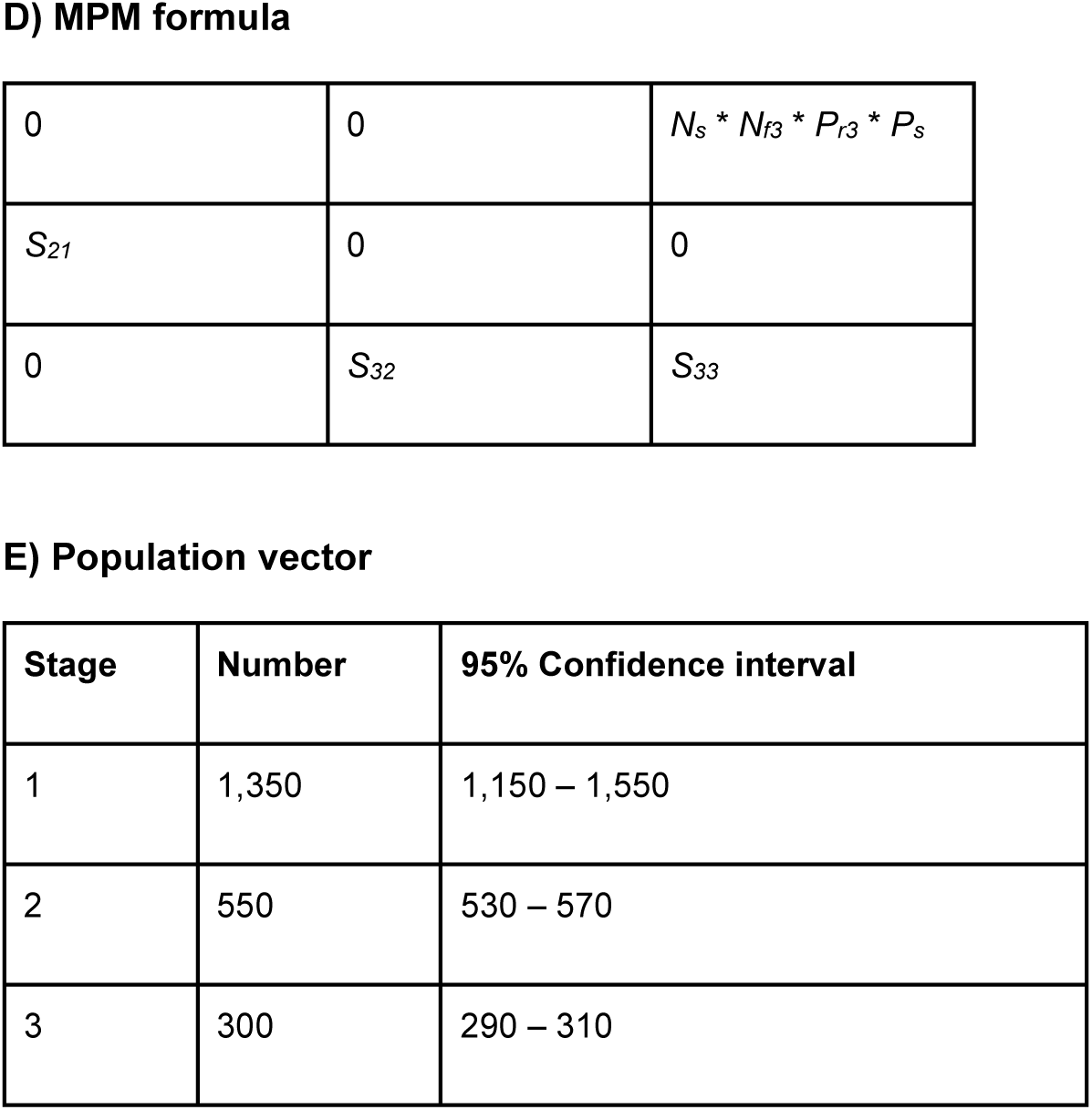

## The theory does not stand still: nonlinearity, environment-dependence and multistate models

MPMs have become a predominant approach in the toolbox of population ecologists partly due to their simplicity of construction and analysis. But the theory underlying matrix-based demography does not stand still, and in the last 20 years it has enlarged dramatically. These new methods produce models whose structure does not fit into the frameworks for reporting that seemed so comprehensive in the past. Furthermore, the body of data available for demography has grown rapidly from long-term ecological research and monitoring networks (*e.g.*, LTERs in USA (Hobbie, Carpenter, Grimm, Gosz, & Seastedt, 2003), Wytham Woods (Macdonald, Newman, Stewart, Domingo-Roura, & Johnson, 2002; Simmonds, Cole, Sheldon, & Coulson, 2020)) and remote sensing approaches (*e.g.*, tagging, drone-monitoring projects (Cubaynes et al., 2022)). These datasets, and recent advances in MPM theory and methods, enable researchers to link population dynamics and demography to environmental conditions and multiple individual traits (*e.g.*, sex and age (Childs, Sheldon, & Rees, 2016); age and kinship (Caswell, 2019b, 2020)) rather than a single trait. These advances offer benefits for the study of population responses to extreme climate (Jenouvrier et al., 2022), as well as more nuanced investigations of comparative and evolutionary demography (Childs et al., 2016). In turn, in this section, we overview some exciting areas of structured demography that can open novel research questions for the modern demographer and list some of the challenges they pose for communication and reporting.

### Non-linear dynamics

Nonlinear MPMs are those in which entries of the projection matrix depend on the population state (numbers and structure) and may be frequency- or density-dependent. Frequency-dependent nonlinearities depend only on the relative abundance of stages; they occur in two-sex models in which mating depends on the relative abundance of males and females, and in population genetic models where dynamics depend on the relative abundance of genotypes (de Vries & Caswell, 2019). Density-dependent models depend on the abundance and structure of the population; recent examples include Pardini et al. (2009) and Shyu et al. (2013) for analyses of control strategies for garlic mustard (Alliaria petiolata) and de Vries et al. (2020) for laboratory studies of pesticide resistance in Tribolium.

The analysis of nonlinear MPMs focuses on demographic outcomes different from those of linear models; equilibria, attractors, bifurcations, oscillations, and stability (see Caswell 2001, Chapters 16 and 17). However, what makes these models problematic for the current status of COMPADRE and COMADRE is that the unit of the model is not a matrix, but rather a matrix function, in which the entries of the projection matrix are functions of the state of the population. Sensitivity analyses are available to study pretty much any demographic outcome in response to any parameter (Caswell, 2019a), but reporting the functions that define the MPM is not at all standardized.

### Environment-dependence

A similar problem arises in environment-dependent MPMs. In such models, some or all of the demographic rates are functions of some aspects of the environment; *e.g.*, polar bears as functions of statistics of Arctic sea ice (Regehr, Hunter, Caswell, Amstrup, & Stirling, 2010), sifaka as functions of rainfall (Lawler et al., 2009), the emperor penguin as a function of seasonal sea ice patterns in the Antarctic (Jenouvrier et al., 2012), and the North Atlantic right whale as functions of time and of trends in time (Fujiwara & Caswell, 2001). As with nonlinear MPMs, the model is not a matrix, but a function that maps from the environmental variable(s) to the entries in the matrix. Protocols for reporting such functions are not yet available but are important to develop.

### Multi-state models

An exciting emerging area of demographic research is the construction and analysis of multistate MPMs, in which individuals are classified by more than one state variable. This includes age and stage (Caswell, 2012; Caswell & Salguero-Gómez, 2013), stage and spatial location (Hunter & Caswell, 2005), stage and genotype (de Vries & Caswell, 2019), stage and infection status (Klepac & Caswell, 2011), age and unmeasured heterogeneity (Hartemink, Missov, & Caswell, 2017), and stage-specific incidence of disease (Caswell & Van Daalen, 2021). A detailed presentation of the methods is given in (Caswell, de Vries, Hartemink, Roth, & van Daalen, 2018) and the extension to more than two state axes (so-called hyperstate matrices) is given in (Roth & Caswell, 2016). The incorporation of additional states enables researchers to tease apart various sources of individual heterogeneity and to ask deeper comparative and evolutionary questions. For example, maternal age has a strong impact on vital rates in monogonont rotifers (Bock, Jarvis, Corey, Stone, & Gribble, 2019). Applying vec-permutation methods (Caswell, 2012) to build multi-state MPMs has allowed researchers to quantify the population-level impacts of the observed maternal age effect and to investigate the evolutionary processes that can lead to this type of senescence in rotifers (Hernández et al., 2020). Multi-dimensional MPMs and Markov chain approaches (van Daalen & Caswell, 2017) have been particularly important in the study of “luck” in life histories, which explores why some individuals live long and prosper, while others do not (Snyder & Ellner, 2018). In studies of “luck,” variance among individuals for a life history outcome is partitioned into contributions from between-group and within-group variation (*e.g.*, van Daalen & Caswell, 2017; Snyder & Ellner, 2018). Examples of sources of individual heterogeneity include maternal age (van Daalen, Hernández, Caswell, Neubert, & Gribble, 2022), birth-year environment (Snyder & Ellner, 2022), and genetic variation (Steiner, Tuljapurkar, & Roach, 2021). The within-group variation is called individual stochasticity or “luck” and arises from the fact that vital rates are probabilistic processes.

These models pose a challenge for reporting because the MPM consists not of a single matrix, but of four sets of matrices. Consider an age x stage-classified model. It is composed of a set of matrices giving transitions among stages for each age class, a set of matrices ***D*** giving age transitions for each stage, a set of matrices ***F*** giving stage-specific fertility for each age, and a set of matrices ***H*** that allocate newborn offspring to the appropriate ages. These are assembled into block structured transition and fertility matrices from which all the usual demographic outcomes can be calculated and related to both age and stage (*e.g.*, see Caswell and Salguero-Gómez (2013) for an analysis of selection gradients for both age and size).

### Communication

The reproducibility of this new generation of MPMs that include more realism and accommodate a higher degree of individual heterogeneity requires more metadata than in previous, simpler MPM examples. In cases where the matrix elements and/or underlying vital rates are formulas, the full formulas, clearly indicating the variables that shape them, and the domain of values that they can occupy is necessary. In the case of multi-stage MPMs, it is necessary to know both the vital rates themselves, as well as the matrix construction procedure (for example, Hernández et al. (2020) uses a stage-within-age construction). For MPMs that incorporate a Markov chain with rewards, the definitions of absorbing states and rewards must be clearly presented; reproducibility also requires knowing the mixing distribution that was used for variance analysis (see van Daalen et al., 2022). The development of MPMs to accommodate these richer realities requires a more detailed report of their data and metadata.

## Discussion

Demographic research has come a long way since the introduction of age-based (Leslie, 1945) and stage-based matrix models (Lefkovitch, 1965). Advances in this field have been fuelled partly by clear communication of methods and associated code. We aim to continue this expansion with MPM communication.

As the depth and breadth of the literature continues to expand, we are starting to build a comprehensive picture of demography across the spectrum of life (Adler et al., 2014; McDonald et al., 2016; Paniw et al., 2018, 2019; Jackson et al., 2022; Le Coeur et al., 2022). Through the work of the COMPADRE and COMADRE databases, we have come to appreciate the utility and opportunities of a standardised way of compiling MPMs. Indeed, a significant portion of the time (>50%) we spend curating these databases is actually not on digitising, error-checking, and complementing data, but on contacting authors for clarification and request of missing data and metadata. Through this arduous process, we have identified valuable-yet typically missing-information in MPMs. Whilst the missing data highlighted here as being particularly important primarily reflects the interests and perspectives of comparative demographers, including the data outlined in the standardised method would benefit demography as a whole.

This paper intends to act as a useful reference for authors, editors, reviewers, managers/conservationists and comparative demographers. Furthermore, we hope this manuscript will promote a constructive discussion on the purpose, construction, and presentation of stage-based demographic information. Box 1 contains a comprehensive example of the key information we believe should be incorporated into the publication of any MPM. Should the methods suggested here be adopted, there will be clear benefits for the growth of the COMPADRE and COMADRE demographic databases; however, we believe these benefits extend beyond COMPADRE and COMADRE users toward the whole field of population ecology. A greater level of detail and transparency when describing how and why an MPM is produced will result in greater accuracy, accessibility, reproducibility and citability – this has clear benefits to the field as a whole and to individual researchers. In addition, greater consistency and transparency facilitates peer review, and indeed these guidelines may offer a tool that can be cited by associate editors and peer-reviewers who may frequently advocate some (or all) of the steps suggested herein. Furthermore, adoption of the steps suggested here may increase confidence in the results presented and facilitate learning/uptake of MPMs by early career researchers.

Finally, we close with a caution. We have used the term “accurate” at points throughout this paper, applied to MPMs, but we must acknowledge that there is no such thing as an accurate model, be it an MPM or any other type. A model is a series of choices, choices of aspects that are included and aspects that are neglected. Model selection techniques such as AIC (Anderson & Burnham, 2002) make these choices explicit and measure their support in terms of likelihood. But even without using the explicit statistical method, the message is clear. Choices of *i*-state variables, of projection intervals, of types of time variation, of functional dependence on a chosen set of environmental factors, … all of these are inaccurate. The point is not to seek for accuracy, it is to be clear about communicating the choices you made in constructing the model, the analyses you chose to apply and the interpretation of the results. An “accurate” model of an ecological system, experimental or observational, would be as complicated as the real system. That does not end well (Borges, 1998).

## Conclusion

Open data in ecology is a growing movement (Nadrowski et al., 2013; Gallagher et al., 2020; Salguero-Gómez et al., 2021). As we move ever further into the data-driven era of quantitative biology, data standards become increasingly important (Gallagher et al., 2020) Other fields have jumped on this standardisation bandwagon (*e.g.*, meta-analyses (PRISMA method: Moher et al., 2009, 2015), genome/amino acid sequences (FASTA format: *e.g.*, Pearson, 2016) and qPCR detection limits (Forootan et al., 2017)). It is time population ecologists join that movement. MPMs are a mathematical representation of discrete transitions and per-capita contributions of individuals in a population across discrete time steps. To fully harness the immense collective potential of the thousands of MPMs produced to date, we need thorough data archiving, community derived standards and biological nuance. Whilst the nuance for a fruit fly population would be inherently different from a fruit tree population, our proposed standardisation protocol ensures future insights, connections and impact will grow due to the effective communication of MPMs.

## Author contributions

This paper was conceptualized at a workshop hosted by JM, DB and DH. Subsequently, SJLG, SR, CMH and RS-G generated the first draft of the manuscript with ideas contributed from all authors. SJLG conducted the survey. SJLG, DS, NN and ASSP conducted the screen of papers from COMPADRE and COMADRE.

## Acknowledgements

We thank the hundreds of population ecologists who have contributed open-access matrix population models ready for fully reproducible research, and those who have, throughout the last 15 years, answered our emails asking for additional data and metadata. Conversations leading to this manuscript were initiated during a workshop held at the University of Exeter Cornwall campus with support of NERC grant (NE/N006798/1) to JM and DH. We acknowledge the support of the Evolutionary Demography Laboratory at the Max Planck Institute for Demographic Research (MPIDR) in the development of the COMPADRE Plant Matrix Database and COMADRE Animal Matrix Database, and the maintenance of COMPADRE & COMADRE through the distributed network of digitising nodes including MPIDR, University of Oxford, University of Exeter, Southampton University, Trinity College Dublin, Lincoln Park Zoo, University of Southern Denmark, and iDiv. RS-G was supported by a NERC Independent Research Fellowship (NE/M018458/1). CMH was supported by a US-NSF grant (DEB-1933497). PC was supported by a Maria Zambrano Next Generation EU Fellowship. JC-C, ORJ, RS-G, and CCT were supported by an NSF Advances in Bioinformatics Development Award (#DBI-1661342). DB was funded by NERC Discovery Science grant NE/L007770/1. TK was supported by the Helmholtz Association and the Alexander von Humboldt foundation. And lastly, this paper is in memoriam of our dear friend and colleague James W. Vaupel, who sadly passed before the submission of this manuscript. His multiple contributions to demography will no doubt outlive multiple Bristlecone pine generation times.

## Conflict of interest

The authors declare no conflict of interest.

## SUPPLEMENTARY ONLINE MATERIALS

S1: A survey on matrix communication

Title: MPMs in the Literature

Section 1: Description

Thank you for taking this 2-minute survey. We really value your input. The main goal of this survey is to assess the ability of publications using MPMs to impart sufficient information about MPM construction.

All answers are anonymous, and participants can end the survey at any time.

Below, you will find two questions gauging your use of matrix population models (MPMs) in your research. Subsequently, there are statements about how MPMs and their associated information are reported in the literature. To complete the survey, please indicate to what extent you agree with the proposed statements.

The ratings represent:

1 = strongly disagree

2 = disagree

3 = neither agree nor disagree

4 = agree

5 = strongly agree

Section 2: General information

1. How many years have MPMs been involved in your research

a. Possible answers:

i. 1
ii. 2
iii. 3
iv. 4
v. 5
vi. 5+
2. You are comfortable at building stage-structured population models (*i.e.*, MPMs or IPMs).

a. Possible answers:

i. Strongly disagree
ii. Disagree
iii. Neither agree nor disagree
iv. Agree
v. Strongly agree

Section 3: MPMs in research

1. Population ecologists who use MPMs describe the methods associated with building the MPMs form raw data adequately for reproducibility in peer-reviewed publications.

a. Possible answers:

i. Strongly disagree
ii. Disagree
iii. Neither agree nor disagree
iv. Agree
v. Strongly agree
2. Papers using MPMs identify and clearly define the names of stage/age/size classes used in their structure.

a. Possible answers:

i. Strongly disagree
ii. Disagree
iii. Neither agree nor disagree
iv. Agree
v. Strongly agree
3. Census duration, the period of time data are recorded (*e.g.*, 6 months, 10 years), is reported with the MPMs in peer-reviewed publications.

a. Possible answers:

i. Strongly disagree
ii. Disagree
iii. Neither agree nor disagree
iv. Agree
v. Strongly agree
4. Projection interval, the period of time between censuses (*e.g.*, 3 months, 1 year), is reported with the MPMs in peer-reviewed publications.

a. Possible answers:

i. Strongly disagree
ii. Disagree
iii. Neither agree nor disagree
iv. Agree
v. Strongly agree
5. Authors detail the formulas used to calculate vital rates (*i.e.*, attributions from survival, sexual reproduction, clonal reproduction, retrogression towards a single matrix element) in peer-reviewed publications.

a. Possible answers:

i. Strongly disagree
ii. Disagree
iii. Neither agree nor disagree
iv. Agree
v. Strongly agree
6. The life-cycle graph, which illustrates the discrete transitions involved in an MPM, is present in peer-reviewed publications using MPMs.

a. Possible answers:

i. Strongly disagree
ii. Disagree
iii. Neither agree nor disagree
iv. Agree
v. Strongly agree
7. Population vectors, which define the structure of the population, are explicitly reported in peer-reviewed publications using MPMs.

a. Possible answers:

i. Strongly disagree
ii. Disagree
iii. Neither agree nor disagree
iv. Agree
v. Strongly agree
8. Papers using MPMs display the MPM in a table or in the supplementary information.

a. Possible answers:

i. Strongly disagree
ii. Disagree
iii. Neither agree nor disagree
iv. Agree
v. Strongly agree
9. Given the current state of peer-reviewed publication practice around MPMs, I think a standardized method of MPM reporting is necessary for the coherent communication of MPMs in the literature.

a. Possible answers:

i. Strongly disagree
ii. Disagree
iii. Neither agree nor disagree
iv. Agree
v. Strongly agree

## Notes

### Competing Interest Statement

The authors have declared no competing interest.

